# Cis- and Trans-Acting Expression Quantitative Trait Loci Differentially Regulate Gamma-Globin Gene Expression

**DOI:** 10.1101/304899

**Authors:** Elmutaz M. Shaikho, John J. Farrell, David H. K. Chui, Paola Sebastiani, Martin H. Steinberg

**Author notes:** Correspondence: Martin H. Steinberg, 72 E. Concord St., Boston, MA 02118.

## Abstract

Genetic association studies have detected two trans-acting quantitative trait loci (QTL) on chromosomes 2, 6 and one cis-acting QTL on chromosome 11 that were associated with fetal hemoglobin (HbF) levels. In these studies, HbF was expressed as a percentage of total hemoglobin or the number of erythrocytes that contain HbF (F-cells). As the *γ*-globin chains of HbF are encoded by two non-allelic genes *(HBG2, HBG1)* that are expressed at different levels we used normalized gene expression and genotype data from The Genotype-Tissue Expression (GTEx)-project to study the effects of cis- and trans-acting HbF expression or eQTL. This allowed us to examine mRNA expression of *HBG2* and HBG1individually. In addition to studying eQTL for globin genes we examined genes co-expressed with *HBG1*, studied upstream regulators of *HBG1* co-expressed genes and performed a correlation analysis between *HBG2* and *HBG1* and known HbF regulators. Our results suggest differential effect of cis and trans-acting QTL on *HBG* and *HBG1* expression. Trans-acting eQTLs have the same magnitude of effect on the expression of both *HBG2* and *HBG1* while the sole cis-acting eQTL affected only *HBG2*. Furthermore, the analysis of upstream regulators and the correlation analysis suggested that *BCL2L1* might be a new potential trans-acting HbF activator. HbF is the major modulator of the phenotype of sickle cell anemia and β thalassemia. Depending on the effect size, modification of trans-acting elements might have a greater impact on HbF levels than cis-acting elements alone.

## Introduction

Genome-wide and other genetic association studies have detected many single nucleotide polymorphisms (SNPs) in chromosomes 2, 6, and 11 that were associated with fetal hemoglobin (HbF) levels in normal individuals and in patients with sickle cell anemia and β thalassemia (Thein et al. 1987; Craig et al. 1996; Garner et al. 1998; Uda et al. 2008; Mtatiro et al. 2014). These studies used either HbF protein levels–expressed as a percentage of total hemoglobin– or the percentage of all erythrocytes that had HbF detected by immunofluorescence (F-cells) as a quantitative trait. HbF is a tetramer composed of two *α* and two *γ*-polypeptide subunits. The *γ*-globin subunits, which characterize HbF, are encoded by two closely linked genes–from 5’ to 3’, *HBG2* and *HBG1* (together, *HBG*). The respective polypeptides of the *γ*-globin genes differ by only a single amino acid in position 136; glycine in the ^G^*γ* chain (HBG2) and alanine in the ^A^*γ* chain (HBG1) (Schroeder et al. 1968). In adults, these genes are expressed at a ratio of 2:3 (Terasawa et al. 1980). Expression quantitative trait loci (eQTLs) are genomic loci that lead to variability in mRNAs expression levels and can be identified by analyzing global RNA expression using the expression levels of genes as quantitative phenotypes. Differential expression of *HBG* in whole blood can be detected by mRNA analysis that may provide an understanding of whether eQTL associated with *HBG* affects the expression of one or both genes.

RNA sequence data (RNA-seq) and whole genome sequences from primary erythroid progenitors expressing *HBG* would provide the ideal data set to study cis and trans-acting elements that regulate HbF expression. However, such data are rarely available publicly. The Genotype-Tissue Expression (GTEx)-project provides an alternative resource to study *HBG* gene expression, the regulatory elements of these genes and the effects of genetic variation of these elements (GTEx_Consortium 2013). GTEx was launched in 2010 to create a publicly available database of genotype and tissue expression. Samples come from either deceased or surgical donor’s organ/tissues. These human tissue samples then undergo RNA-seq and genome-wide SNP analysis. The tissues sampled include peripheral blood, which contains reticulocytes that express hemoglobin genes. Reticulocytes account for about 10% of total hemoglobin synthesis.

We performed a genome-wide association study (GWAS) to detect eQTLs associated with expression of *HBG* in whole blood, other genes of the β-globin (HBB) gene cluster and with known HbF cis- and trans-acting regulators. In addition, we examined genes co-expressed with *HBG1* to discover enriched pathways, studied upstream regulators of HbF co-expressed genes and performed correlation analysis between *HBG* and known HbF regulators. A similar correlation analysis included *BCL2L1*, a potential HbF activator we found in our analysis. We hypothesized that trans-acting elements that are transcription factors affect the expression of both *HBG2* and *HBG1*, while the known cis-acting QTL in the promoter of *HBG2* is likely to be associated with the expression of this gene only.

## Results

### Preliminary studies

GTEx whole blood RNA-seq data detected HbF eQTLs in *BCL11A*, HMIP and in the *HBB* gene cluster using HbF gene expression as a surrogate for HbF protein levels, validating the utility of this data set for examining HbF-associated eQTL (**Table 1 and Fig. 1A-F**).

**Table 1.** **SNPs from previous GWAS and their association with *HBG1* and *HBG2* expression in whole blood samples available in the GTEx portal**.

| Gene Symbol | Variant Id | SNP | P-Value | Effect Size | T-Statistic | Standard Error |
| --- | --- | --- | --- | --- | --- | --- |
| ***HBG1*** | 2_60718043_T_G_b37 | rs1427407 | 1.20E-11 | -0.46 | -7.1 | 0.065 |
| ***HBG1*** | 11_5263683_C_T_b37 | rs10128556 | 0.89 | -0.0085 | -0.14 | 0.059 |
| ***HBG1*** | 6_135419018_T_C_b37 | rs9399137 | 2.20E-15 | 0.51 | 8.4 | 0.061 |
| ***HBG2*** | 2_60718043_T_G_b37 | rs1427407 | 1.10E-09 | -0.46 | -6.3 | 0.073 |
| ***HBG2*** | 11_5263683_C_T_b37 | rs10128556 | 2.60E-16 | 0.51 | 8.7 | 0.058 |
| ***HBG2*** | 6_135419018_T_C_b37 | rs9399137 | 1.70E-09 | 0.44 | 6.2 | 0.07 |

**Figure 1.**
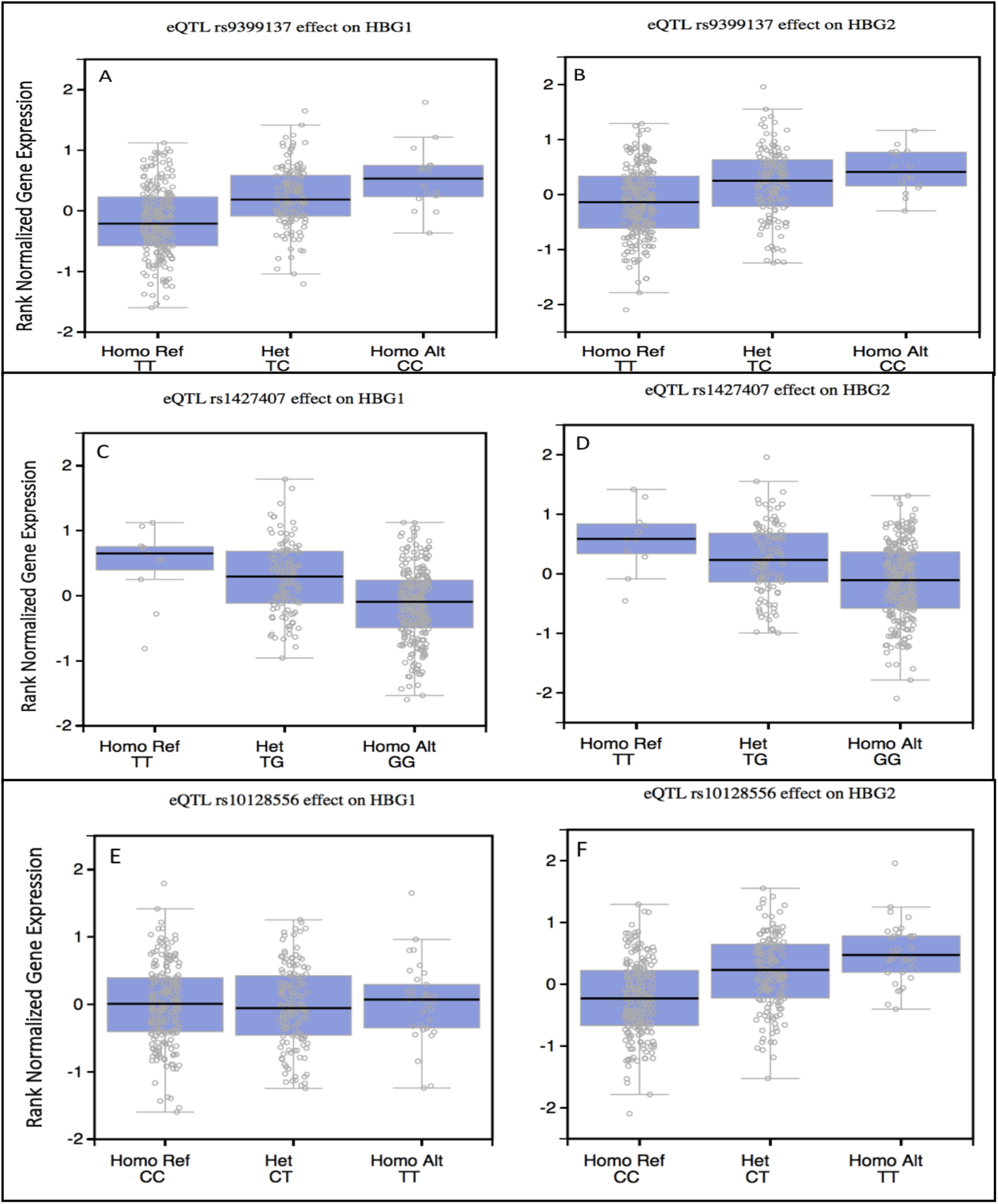
**Effect of rs1427407 (2:60718043_T/G), rs10128556 (11:5263683_C/T), and rs9399137 (6:135418632_TTAC/T) on *HBG1* and *HBG2* expression in GTEx data set version 6.** P-values are above genome-wide significance for rs1427407 (*BCL11A*, chr2) and rs9399137 (*MYB*, chr6) association with expression of both *HBG* genes. Rs10128556 (chr 11) is significantly associated with only *HBG2* (p-value of 2.60E-16) while it has no effect on *HBG1* (p-value 0.89) Het denotes heterozygous, and Homo, homozygous.

### Genome-wide eQTL association analysis

Twenty-seven SNPs were eQTL associated with *HBG1* (p-value ≤ 5 × 10-8). Of these 27, 21 were intergenic variants in the HMIP region on chromosome 6p, five were *BCL11A* variants on chromosome 2p, and one was on chromosome 11p (**Fig. 2A, Table S1**). The SNP rs66650371, the functional 3-bp deletion in the *MYB* enhancer, is one of the most significant SNPs in the HMIP region. The most significant SNP of the five *BCL11A* variants was rs1427407, which in other studies was the functional SNP in the erythroid-specific enhancer of this gene (Bauer et al. 2013).

**Figure 2.**
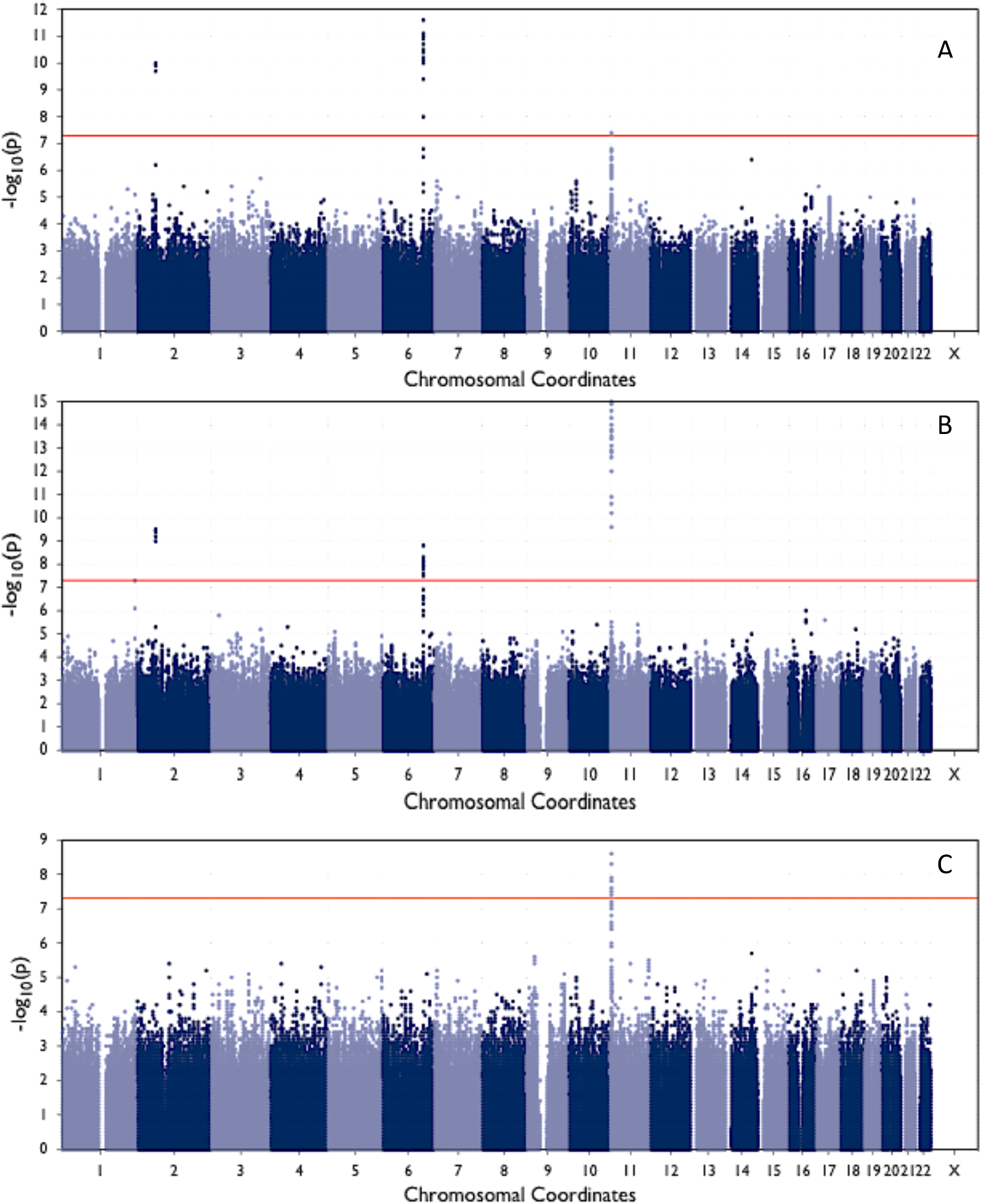
**Manhattan plots for *HBG* eQTLs in 338 whole blood samples. Fig. 2A** shows *HBG1* eQTL; **Fig. 2B** shows *HBG2* eQTL; **Fig. 2C** shows *HBG1* eQTL conditioned on rs66650371, rs1427407 and rs7482144 genotype. The red line indicates genome-wide significance levels

Forty-nine SNPs meeting genome-wide significance levels were eQTL for *HBG2*. Seventeen were intergenic variants in the HMIP region on chromosome 6, five were *BCL11A* variants on chromosome 2p, 26 were on chromosome 11p (**Fig 2B, Table S2**) and one was on chromosome 1. The most significant SNP in chromosome 11 was rs7482144 and this was associated only with the expression of *HBG2*; rs16912979, which is located in hypersensitive site (HS) 4 of the locus control region (LCR) of the *HBB* gene complex, was also associated solely with *HBG2* expression. The SNP rs66650371 *(MYB)* and rs1427407 *(BCL11A)* were significantly associated with both *HBG2* and *HBG1*. The conditional analysis of *HBG1* showed that after adjusting for the effects of rs1427407, rs66650371, and rs7482144, there was no association on chromosome 2 or chromosome 6 (**Fig. 2C, Table S3**).

Conditional analysis of *HBG2* showed that no SNP was significantly associated with *HBG2* expression after adjusting for rs7482144, rs1427407, and rs66650371 (**Fig. S1, Table S4**). There was no SNP with genome-wide significance associated with expression of *BCL11A* and *ZBTB7A* (**Figs. S2, S3; Tables S5, S6**). One SNP on chromosome 6 was associated with *MYB* expression, another on chromosome 6 was associated with *HBD* expression, and one on chromosome 12 was associated with *KLF1* expression (**Figs. S4, S5, S6; Tables S7, S8, S9**). Variants on chromosome 1, 2, 6, 7, 14, and 20 were significantly associated with *HBB* expression (**Fig. S7, Table S10**).

### Association of the erythroid-specific BCL11A enhancer variant rs1427407 with whole blood gene expression

Linear regression analysis showed that the genotype of rs1427407 was significantly associated with *HBG1* and *HBG2* expression in whole blood (**Table 2**) after correction for multiple testing using FDR, p-values were 2.04E-06 and 3.63E-06 for *HBG1* and *HBG2*, respectively. This SNP was not a eQTL for any other gene expressed in peripheral blood.

**Table 2.** **Effect of rs1427407 on whole blood gene expression.** ENS_ID is ensemble gene ID, LogFC is log fold change, aveExpr is average expression, and adj.P.Val is FDR adjusted p-value.

| ENS_ID | Gene Symbol | logFC | AveExpr | P.Value | adj.P.Val |
| --- | --- | --- | --- | --- | --- |
| ENSG00000213934.5 | *HBG1* | 0.502491942 | -4.32E-06 | 8.51E-11 | 2.04E-06 |
| ENSG00000196565.8 | *HBG2* | 0.507932715 | -7.56E-06 | 3.03E-10 | 3.63E-06 |
| ENSG00000187017.10 | *ESPN* | 0.307905604 | -1.45E-07 | 1.80E-05 | 0.143668299 |
| ENSG00000239405.1 | *TMED10P2* | 0.362195583 | 0.002468525 | 7.07E-05 | 0.423972382 |
| ENSG00000010626.10 | *LRRC23* | 0.277170914 | 2.20E-07 | 9.30E-05 | 0.427249648 |
| ENSG00000248309.1 | *MEF2C-AS1* | 0.320432287 | -9.14E-08 | 0.000106933 | 0.427249648 |
| ENSG00000160401.10 | *CFAP157* | -0.342869883 | -8.98E-07 | 0.000164101 | 0.561998024 |
| ENSG00000135414.5 | *GDF11* | -0.163176468 | -3.51E-08 | 0.000286461 | 0.777799594 |
| ENSG00000255318.1 | - | -0.358828046 | 0.000665361 | 0.000316391 | 0.777799594 |

### HBG1 differential co-expression analysis

Twelve genes were differentially co-expressed with *HBG1* after FDR p-value adjustment (**Table S11**). The pathway enrichment analysis for the top100 co-expressed genes did not include any known pathway involved in HbF regulation among the top 10 enriched pathways. Of 452 upstream regulators that regulate the top100 *HBG1* co-expressed genes, 360 regulated *BCL2L1* (**Table S12**). GATA1and *KLF1* were also in top five statistically significant upstream regulators.

### Correlation analysis between known and potential HbF regulators and HBG

The correlation between *HBG1 and HBG2* normalized gene expression and the expression of *BCL11A, KLF1, MYB, ZBTB7A, SIRT1* and *BCL2L1* in the GTEx whole blood data set, showed that *BCL2L1* had correlation coefficients of 0.59 and 0.55 for *HBG1* and *HBG2*, respectively with a p value of 2.2E-16. (**Table 3**).

**Table 3.** **Pearson correlation between *HBG* and known or potential HbF regulators using RNA-seq from 338 whole blood samples from GTEx.**

| Regulators | *HBG1* Correlation | P-value | *HBG2* Correlation | P-value |
| --- | --- | --- | --- | --- |
| *BCL2L1* | 0.59 | 2.20E-16 | 0.55 | 2.20E-16 |
| *KLF1* | 0.47 | 2.20E-16 | 0.44 | 2.20E-16 |
| *MYB* | 0.27 | 2.94E-07 | 0.27 | 2.8E-07 |
| *BCL11A* | 0.15 | 0.003914 | 0.17 | 0.001694 |
| *SIRT1* | -0.06 | 0.238 | -0.06 | 0.29 |
| *ZBTB7A* | -0.0045 | 0.9334 | 0.067 | 0.212 |

In primary human fetal liver proerythroblasts, *BCL2L1, KLF1*, and *SIRT1* were positively correlated with *HBG* expression with correlation coefficients of 0.93, 0.92, and 0.62, respectively; *BCL11A* and *MYB* were negatively correlated with *HBG1* with correlation coefficients of −0.63, and −0.64, respectively; *ZBTB7A* was weakly correlated with *HBG1*, and the p-value for association was not significant (**Table 4**).

**Table 4.** **Pearson correlation between *HBG* and known or potential HbF regulators using RNA-seq primary human fetal liver proerythroblasts.**

| Regulators | *HBG1* Correlation | P-value | *HBG1* Correlation | P-value |
| --- | --- | --- | --- | --- |
| *BCL2L1* | 0.93 | 3.85E-07 | 0.93 | 3.85E-07 |
| *KLF1* | 0.92 | 9.24E-07 | 0.92 | 9.24E-07 |
| *MYB* | -0.64 | 0.01 | -0.63 | 0.01 |
| *BCL11A* | -0.63 | 0.01 | -0.63 | 0.01 |
| *SIRT1* | 0.62 | 0.01 | 0.62 | 0.01 |
| *ZBTB7A* | -0.38 | 0.17 | -0.38 | 0.16 |

## Discussion

Among eQTL significantly associated with *HBG* expression 21 were intergenic variants in HMIP on chromosome 6p. The functional 3-bp deletion (rs66650371) was one of the most significant eQTL in this *MYB* enhancer (Farrell et al. 2011; Stadhouders et al. 2014). The SNP rs66650371 was in strong LD with 20 other SNPs in chromosome 6p in Europeans, South and East Asians, and admixed Americans (**Table S13**). A long non-coding or lncRNA containing the site of rs66650371was transcribed from this enhancer. Its downregulation in an erythroid cell line that expressed adult hemoglobin (HbA) was associated with a 200-fold increase in *HBG* expression and a 20-fold increase in HbF (Morrison et al. 2017). The eQTL at rs66650371 further supports the functional importance of this SNP. Five *BCL11A* SNPs in chromosome 2p were in very strong LD across populations (**Table S14**). The most significant, rs1427407, is a functional erythroid-specific enhancer variant that altered the expression of *BCL11A* (Bauer et al. 2013). Both rs1427407 and rs66650371are trans-acting elements and had the same magnitude of effect on *HBG1* and *HBG2* (**Fig. 1C, 1D; Fig. 3**). These observations are consistent with our hypothesis that trans-acting elements affect expression of both *γ*-globin genes.

**Figure 3.**
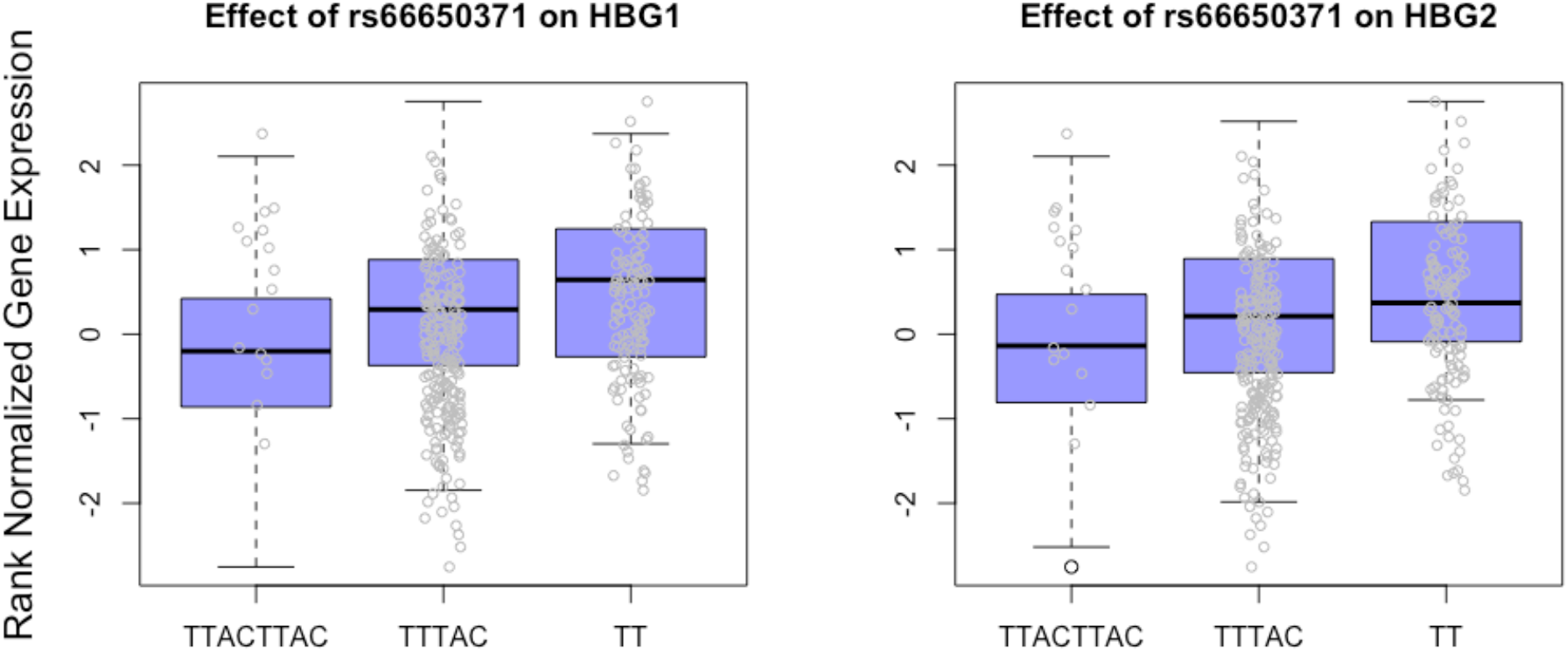
**Effect of rs66650371 (6:135418632_TTAC/T) genotypes on *HBG1* and *HBG2* expression in 338 whole blood samples.** P-values of genome-wide association are 6.49E-11 and 5.86E-09 for *HBG1* and *HBG2*, respectively.

The SNP rs10128556 in the pseudogene *HBBP1* on chromosome 11p was in strong LD with rs7482144 and was reported to have an effect independent of rs7482144 in African American sickle cell disease patients and be the likely functional variant modulating cis-acting HbF gene expression (Galarneau et al. 2010). However, rs7482144 is located within the *HBG2* promoter and highly linked to rs368698783 in *HBG1* promoter. The SNP rs368698783 together with rs7482144 are associated with reduced methylation in six CpG sites flanking the transcription start site of *HBG* (Chen et al. 2017). Moreover, rs7482144 alters a putative binding motif for ZBTB7A, a silencer of *HBG* expression (Masuda et al. 2016; Shaikho et al. 2016).

Strong evidence supporting differential regulation of *HBG* by cis- and trans-acting elements comes from clinical observations of patients with the Arab Indian (AI) and Senegal haplotypes of the sickle hemoglobin (HbS) gene and from the reports of hereditary persistence of HbF (HPFH) caused by point mutations in *HBG* promoters. Patients with the AI and Senegal haplotypes, which are the only *HBB* haplotypes containing rs7482144, had increased levels of HbF that was predominantly of the ^G^*γ*-globin type (Nagel et al. 1985; Ballas et al. 1991; Rahimi et al. 2015). Many mutations have been described in the promoters of *HBG2* and *HBG1* that cause the phenotype of HPFH. For each of these mutations, depending on the affected gene, either HBG2 or HBG1usually comprises more than 90% of total g globin (Wood 2001). The T-C mutations at −175 and −173 in the promoter of *HBG1* reactivated ^A^*γ*-globin gene expression in adult erythrocytes of transgenic mice while altering the binding of GATA-1 and Oct-1 and (Liu et al. 2005). Moreover, a C-T polymorphism that is identical to rs7482144 but at position −158 relative to *HBG1* affected *HBG1* expression (Patrinos et al. 1998).

These results suggest that rs7482144 rather than rs10128556 is either the functional cis-acting element regulating *HBG2* expression or the best surrogate for this element. In addition, rs16912979 in HS-4 of the LCR is also solely associated with *HBG2* further supporting the effect of cis-acting elements on a single *γ*-globin gene (**Fig. 4, 5**). This SNP is a member of the 3-SNP T/A/T haplotype (rs16912979, rs7482144, rs10128556) that is exclusive to the AI haplotype and is associated with their high HbF (Vathipadiekal et al. 2016). The T allele of rs16912979 tags a predicted binding site for runt-related transcription factor 1 (RUNX1) in the palindromic region of 5’ HS-4. RUNX1 plays an important role in hematopoiesis and electromobility shift assays (EMSA) suggested an allele-specific binding of RUNX1 to 5’ HS-4 (Dehghani et al. 2016).

**Figure 4.**
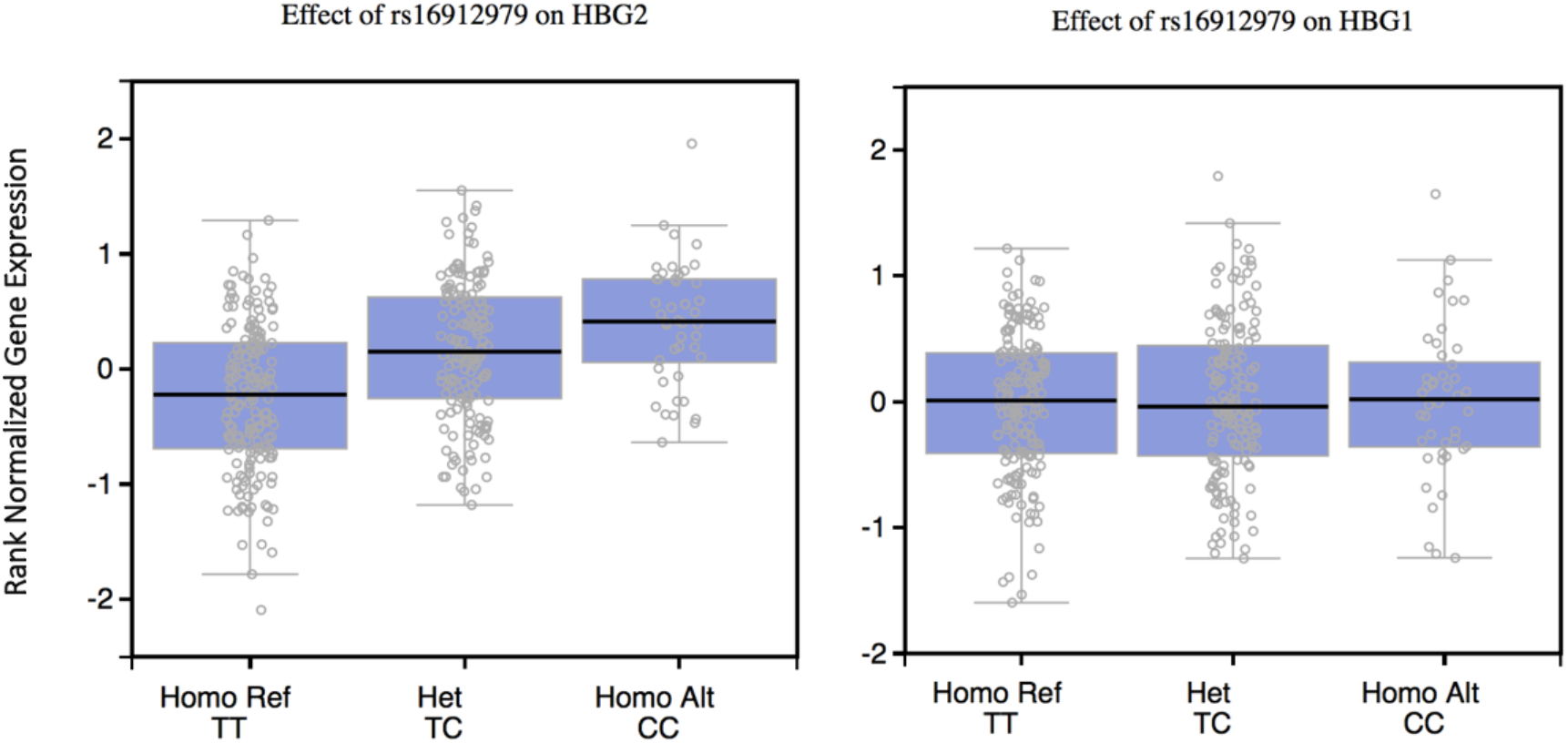
**Effect of rs16912979 (11_5309695_T_C) on *HBG2* and *HBG1* expression in GTEx data set.** p-values of genome wide significance are 7.0e-14 and 0.77, respectively. Het denotes heterozygous, and Homo, homozygous.

**Figure 5.**
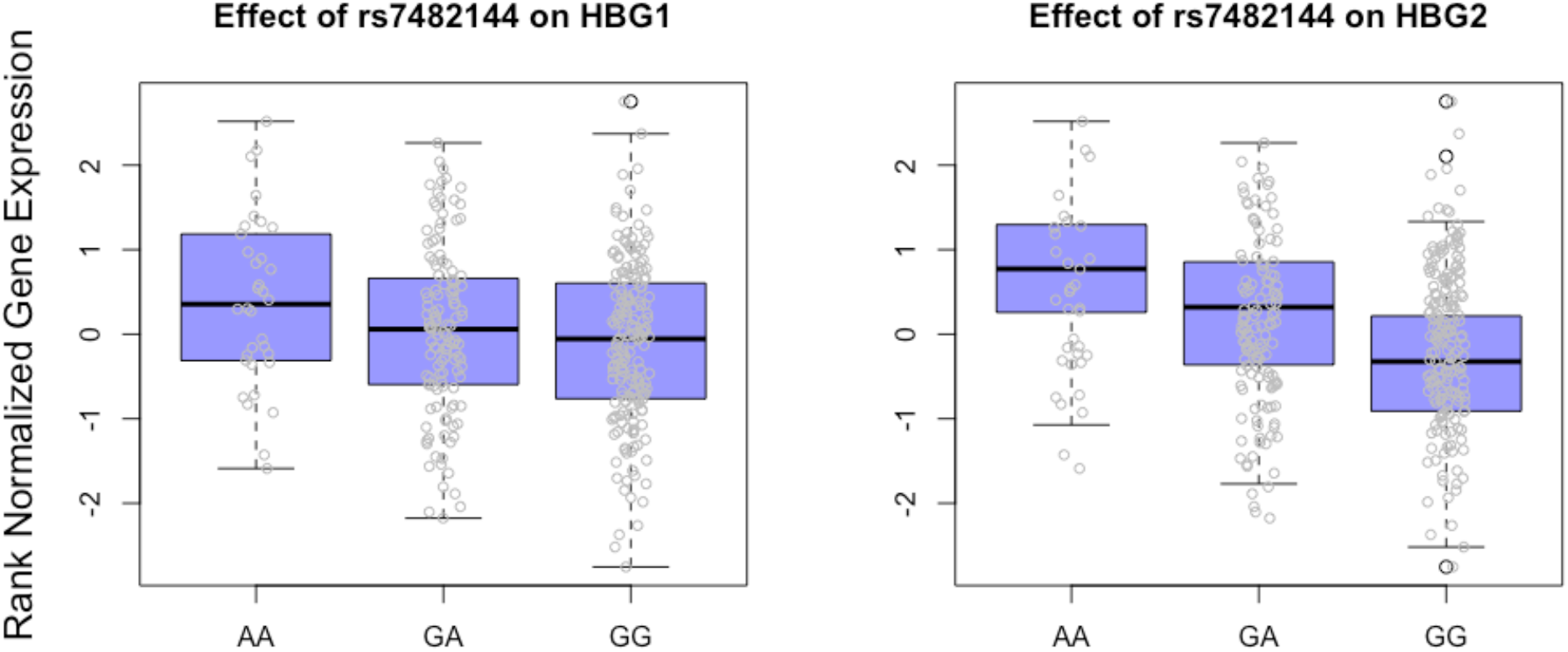
**Effect of rs7482144 (11:5276169_G/A) genotypes on *HBG1* and *HBG2* expression in 338 whole blood samples.** P-values of genome-wide association are 0.038593 and 9.49E-16 for *HBG1* and *HBG2*, respectively.

No common variants affected the expression of *BCL11A*. The *BCL11A* hypersensitive site variant rs1427407 is erythroid specific and in whole blood samples would impact *BCL11A* expression only in reticulocytes. However, *BCL11A* is expressed in leukocytes that are abundant in the blood. *BCL11A* expression in these nucleated cells with their greater transcriptional activity than reticulocytes would render an erythroid-specific signal impossible to detect. Our inability to find genes known to influence *HBG1* in the co-expression analysis and a lack of correlation of known HbF regulators with *HBG* in GTEx data despite a strong relationship in fetal liver proerythroblasts might have a similar explanation and illustrate the main limitation of using GTEx data where globin synthesizing reticulocytes were likely to be <1% of all red blood cells (Means RT 2009). Although few in number, reticulocytes are the only cells in the blood expressing globin genes, and RNA-seq provides sufficient data for most statistical analysis.

The unexpected correlation between *BCL2L1* and *HBG* expression (r 0.58 & 0.55; p=2.30E-16) was validated in fetal proerythroblasts (r> 0.9). *BCL2L1* is a member of bcl-2 gene family involved in anti-apoptotic activities with an important role in erythropoiesis. Induction of bcl-x_L_ *(BCL2L1)* expression by erythropoietin and GATA1 was critical for the survival of late proerythroblasts and early normoblasts (Gregory et al. 1999).The upstream regulation analysis shows that *BCL2L1* was regulated by 80 percent of the regulators that regulate the top 100 co-expressed genes. *GATA1* and *KLF13* are among these upstream regulators. These data and the correlation of *BCL2L1* with *HBG* suggest a role for *BCL2L1* in HbF regulation. Experimental validation to confirm the relationship between *BCL2L1* and *HBG* and to pinpoint the mechanism through which *BCL2L1* regulates *HBG* expression could lead to the discovery of potential new drug targets or molecules that can be used to induce HbF in sickle cell anemia and β thalassemia.

Cis- and trans-acting regulators have a differential effect on *HBG1* and *HBG2* expression. The trans-acting eQTLs of *BCL11A* and *MYB* enhancers affect expression of both *γ*-globin genes. In contrast, only a single *γ*-globin gene is affected by the cis-acting eQTL. HbF is the major modulator of the severity of both sickle cell anemia and β thalassemia and considerable effort is being made to develop methods to increase *HBG* expression for therapeutic purposes (Deng et al. 2014; Lettre and Bauer 2016). Although clinical observations suggest that an increase of HBG2 alone has some benefit (Ballas et al. 1991; Teixeira et al. 2003; Alsultan et al. 2014), modification of trans-acting elements that affect both HbF genes might have a greater effect on HbF levels and help force a more pancellular distribution that could be therapeutically important (Sankaran et al. 2011; Steinberg et al. 2014; Guda et al. 2015; Masuda et al. 2016).

## Methods

### Genome-wide eQTL association analysis

Whole blood normalized RNA-seq data, 1000 Genome imputed genotypes, and the covariates file of 338 donors were downloaded from the GTEx portal version 6. The covariates used in the original GTEx data analysis included 3 genome-wide genotype principal components (PCs), 35 Probabilistic Estimation of Expression Residuals (PEER) factors to be used as confounders, genotyping platform (Illumina HiSeq 2000 or HiSeq X), and sex (Consortium 2017). We performed genome-wide eQTL analysis using Efficient and Parallelizable Association Container Toolbox (EPACTS) (EPACTS 2017) and selected SNPs with minor allele frequency (MAF) ≥ 0.01 and imputation quality score (R^2) ≥ 0.4 to reduce the rate of false association. We used the first 3 PCs to adjust for population substructure. We also used sex and platform as covariates to remove any bias introduced by these two components. The 35 PEER factors were used to adjust for batch effects and experimental confounders. This standard set of covariates were used in the models to detect eQTLs for *HBG1, HBG2, BCL11A, KLF1, MYB, HBB*, and *HBD*. To detect any significant association after adjusting for the most significant SNPs on chromosomes 2, 6, and 11, we tested association of normalized expression of *HBG1* and *HBG2* conditioned on the genotypes of rs7482144, rs1427407, and rs66650371. Only SNPs that reached genome-wide significance (p-value ≤ 5 × 10^−8^) were considered statistically significant.

### Preliminary studies

To reproduce the genetic association of known QTL variants with *HBG* expression using GTEx RNA-seq data from reticulocytes, three SNPs were selected based on their prior association with HbF levels in GWAS. We chose rs1427407 in the *BCL11A* erythroid-specific enhancer on chromosome 2p (Bauer et al. 2013), rs10128556, which is in high linkage disequilibrium (LD) with the Xmn1 restriction polymorphism (rs7482144) on chromosome 11p (Galarneau et al. 2010), and rs9399137 that is in perfect LD with the 3-base pair deletion (rs66650371) in the *HBS1L-MYB* intergenic polymorphism (HMIP) region on chromosome 6q (Farrell et al. 2011). SNPs in LD with rs66650371and rs7482144 were chosen because GTEx pre-calculated eQTL did not contain these SNPs. The GTEx eQTL calculation tool was used to validate these associations in 338 whole blood samples available in the GTEx portal. We sought to validate the effect of a *BCL11A* erythroid-specific-enhancer SNP on *HBG2* and *HBG1* to confirm that genotype can be used to predict phenotype and vice versa.

### Association of the erythroid-specific BCL11A enhancer variant rs1427407 with whole blood gene expression

We extracted the genotype of rs1427407 form GTEx data set version 6 and used these data to predict the impact of this SNP on global gene expression. The same covariates were used in the eQTL analysis, and Linear Models for Microarray and RNA-seq Data (limma) were used to perform linear regression (Ritchie et al. 2015). We used false discovery rate (FDR) to correct for multiple hypothesis testing.

### HBG1 differential co-expression analysis

Since *HBG2* expression is significantly associated with the genotype of rs7482144, we used *HBG1* expression as a marker of expression of both *γ*-globin genes. A gene expression matrix of normalized log expression of RNA-seq along with covariates including 35 PEER factors and sex from 338 whole blood samples were used as input for limma to perform differential analysis. We used FDR to correct for multiple testing. Using Ingenuity Pathways Analysis (QIAGEN Inc., https://www.qiagenbioinformatics.com/products/ingenuity-pathway-analysis/) we performed pathway analysis using the top100 co-expressed genes to see whether there was any enrichment in pathways involved in HbF regulation. In addition, we examined upstream regulators of the genes in the top100 *HBG1* co-expressed gene list to identify HbF transcriptional regulators that can explain the observed gene expression changes.

### Correlation analysis between known and potential HbF regulators and HBG expression

We performed Pearson correlation between *HBG* expression and the expression of *BCL11A, KLF1, MYB, ZBTB7A, SIRT1* and *BCL2L1* on 338 GTEx whole blood samples. These genes were selected because of their role or putative role as modulators of *HBG* expression, except for *BCL2L1* which was selected based on our *HBG1* co-expression and upstream regulators analysis (Jiang et al. 2006; Zhou et al. 2010; Masuda et al. 2016; Dai et al. 2017).

To replicate parts of our GTEx-based analysis, the same correlation analysis was done on data from erythroid samples acquired from Gene Expression Omnibus (GEO; accession number GSE59089) (Xu et al. 2015). These data contained RNA-seq derived from primary human fetal liver proerythroblasts studied at different developmental stages after shRNA-mediated knockdown of Polycomb Repressive Complex 2 (PRC2) core subunits.

## Data Access

Whole blood normalized RNA-seq data, 1000 Genome imputed genotypes, and the covariates file of 338 donors were downloaded from the GTEx portal version 6.

RNA-seq data from 15 erythroid samples were acquired from Gene Expression Omnibus (GEO; accession number GSE59089).

## Acknowledgments

Funded in part by R01 HL 068970, RC2 HL 101212, R01 87681 (MHS), T32 HL007501 (EMS), from the NIH Bethesda, MD.

## Contribution

PS, JJF, DHKC, MHS supervised the research and edited the paper, EMS conceived the the research, performed the analysis, and wrote the paper.

## References

Alsultan A, Alabdulaali MK, Griffin PJ, Alsuliman AM, Ghabbour HA, Sebastiani P, Albuali WH, Al-Ali AK, Chui DH, Steinberg MH. 2014. Sickle cell disease in Saudi Arabia: the phenotype in adults with the Arab-Indian haplotype is not benign. British Journal of Haematology 164: 597-604.

Ballas SK, Talacki CA, Adachi K, Schwartz E, Surrey S, Rappaport E. 1991. The Xmn I site (−158, C----T) 5’ to the G gamma gene: correlation with the Senegalese haplotype and G gamma globin expression. Hemoglobin 15: 393-405.

Bauer DE, Kamran SC, Lessard S, Xu J, Fujiwara Y, Lin C, Shao Z, Canver MC, Smith EC, Pinello L et al. 2013. An erythroid enhancer of BCL11A subject to genetic variation determines fetal hemoglobin level. Science 342: 253-257.

Chen D, Zuo Y, Zhang X, Ye Y, Bao X, Huang H, Tepakhan W, Wang L, Ju J, Chen G et al. 2017. A Genetic Variant Ameliorates beta-Thalassemia Severity by Epigenetic-Mediated Elevation of Human Fetal Hemoglobin Expression. American Journal of Human Genetics 101: 130-138.

Consortium GT. 2017. Genetic effects on gene expression across human tissues. Nature 550: 204.

Craig JE, Rochette J, Fisher CA, Weatherall DJ, Marc S, Lathrop GM, Demenais F, Thein S. 1996. Dissecting the loci controlling fetal haemoglobin production on chromosomes 11p and 6q by the regressive approach. Nature Genetics 12: 58-64.

Dai Y, Chen T, Ijaz H, Cho EH, Steinberg MH. 2017. SIRT1 activates the expression of fetal hemoglobin genes. American Journal of Hematology 92: 1177-1186.

Dehghani H, Ghobakhloo S, Neishabury M. 2016. Electromobility Shift Assay Reveals Evidence in Favor of Allele-Specific Binding of RUNX1 to the 5’ Hypersensitive Site 4-Locus Control Region. Hemoglobin 40: 236-239.

Deng W, Rupon JW, Krivega I, Breda L, Motta I, Jahn KS, Reik A, Gregory PD, Rivella S, Dean A et al. 2014. Reactivation of developmentally silenced globin genes by forced chromatin looping. Cell 158: 849-860.

EPACTS. 2017. EPACTS: Efficient and Parallelizable Association Container Toolbox. In May 21.

Farrell JJ, Sherva RM, Chen ZY, Luo HY, Chu BF, Ha SY, Li CK, Lee AC, Li RC, Yuen HL et al. 2011. A 3-bp deletion in the HBS1L-MYB intergenic region on chromosome 6q23 is associated with HbF expression. Blood 117: 4935-4945.

Galarneau G, Palmer CD, Sankaran VG, Orkin SH, Hirschhorn JN, Lettre G. 2010. Fine-mapping at three loci known to affect fetal hemoglobin levels explains additional genetic variation. Nature Genetics 42: 1049-1051.

Garner C, Mitchell J, Hatzis T, Reittie J, Farrall M, Thein SL. 1998. Haplotype mapping of a major quantitative-trait locus for fetal hemoglobin production, on chromosome 6q23. American Journal of Human Genetics 62: 1468-1474.

Gregory T, Yu C, Ma A, Orkin SH, Blobel GA, Weiss MJ. 1999. GATA-1 and Erythropoietin Cooperate to Promote Erythroid Cell Survival by Regulating bcl-xL Expression. Blood 94: 87-96.

GTEx_Consortium. 2013. The Genotype-Tissue Expression (GTEx) project. Nature Genetics 45: 580-585.

Guda S, Brendel C, Renella R, Du P, Bauer DE, Canver MC, Grenier JK, Grimson AW, Kamran SC, Thornton J et al. 2015. miRNA-embedded shRNAs for Lineage-specific BCL11A Knockdown and Hemoglobin F Induction. Molecular Therapy 23: 1465-1474.

Jiang J, Best S, Menzel S, Silver N, Lai MI, Surdulescu GL, Spector TD, Thein SL. 2006. cMYB is involved in the regulation of fetal hemoglobin production in adults. Blood 108: 1077-1083.

Lettre G, Bauer DE. 2016. Fetal haemoglobin in sickle-cell disease: from genetic epidemiology to new therapeutic strategies. The Lancet 387: 2554-2564.

Liu LR, Du ZW, Zhao HL, Liu XL, Huang XD, Shen J, Ju LM, Fang FD, Zhang JW. 2005. T to C substitution at −175 or −173 of the gamma-globin promoter affects GATA-1 and Oct-1 binding in vitro differently but can independently reproduce the hereditary persistence of fetal hemoglobin phenotype in transgenic mice. Journal of Biological Chemistry 280: 7452-7459.

Masuda T, Wang X, Maeda M, Canver MC, Sher F, Funnell AP, Fisher C, Suciu M, Martyn GE, Norton LJ et al. 2016. Transcription factors LRF and BCL11A independently repress expression of fetal hemoglobin. Science 351: 285-289.

Means RT GB, General Considerations., Greer JP, Foerster J, Rodgers GM, eds. 2009. Anemia. In Wintrobe’s Clinical Hematology, Vol 1. Lippincott Williams and Wilkins, Philadelphia, PA.

Morrison TA, Wilcox I, Luo HY, Farrell JJ, Kurita R, Nakamura Y, Murphy GJ, Cui S, Steinberg MH, Chui DHK. 2017. A long noncoding RNA from the HBS1L-MYB intergenic region on chr6q23 regulates human fetal hemoglobin expression. Blood Cells, Molecules, and Diseases 69: 1-9.

Mtatiro SN, Singh T, Rooks H, Mgaya J, Mariki H, Soka D, Mmbando B, Msaki E, Kolder I, Thein SL et al. 2014. Genome wide association study of fetal hemoglobin in sickle cell anemia in Tanzania. PloS One 9: e111464.

Nagel RL, Fabry ME, Pagnier J, Zohoun I, Wajcman H, Baudin V, Labie D. 1985. Hematologically and genetically distinct forms of sickle cell anemia in Africa. The Senegal type and the Benin type. New England Journal of Medicine 312: 880-884.

Patrinos GP, Kollia P, Loutradi-Anagnostou A, Loukopoulos D, Papadakis MN. 1998. The Cretan type of non-deletional hereditary persistence of fetal hemoglobin [A gamma-158C-->T] results from two independent gene conversion events. Human Genetics 102: 629-634.

Rahimi Z, Vaisi-Raygani A, Merat A, Haghshenass M, Rezaei M. 2015. Level of Hemoglobin F and Gg Gene Expression in Sickle Cell Disease and Their Association with Haplotype and XmnI Polymorphic Site in South of Iran.

Ritchie ME, Phipson B, Wu D, Hu Y, Law CW, Shi W, Smyth GK. 2015. limma powers differential expression analyses for RNA-sequencing and microarray studies. Nucleic Acids Research 43: e47.

Sankaran VG, Menne TF, Šćepanović D, Vergilio J-A, Ji P, Kim J, Thiru P, Orkin SH, Lander ES, Lodish HF. 2011. MicroRNA-15a and -16-1 act via MYB to elevate fetal hemoglobin expression in human trisomy 13. Proceedings of the National Academy of Sciences doi:10.1073/pnas.1018384108.

Schroeder WA, Huisman TH, Shelton JR, Shelton JB, Kleihauer EF, Dozy AM, Robberson B. 1968. Evidence for multiple structural genes for the gamma chain of human fetal hemoglobin. Proceedings of the National Academy of Sciences of the United States of America 60: 537-544.

Shaikho EM, Habara AH, Alsultan A, Al-Rubaish A, Al-Muhanna F, Naserullah Z, Alsuliman A, Qutub HO, Patra P, Sebastiani P. 2016. Variants of ZBTB7A (LRF) and its β-globin gene cluster binding motifs in sickle cell anemia. Blood cells, molecules & diseases 59: 49.

Stadhouders R, Aktuna S, Thongjuea S, Aghajanirefah A, Pourfarzad F, van Ijcken W, Lenhard B, Rooks H, Best S, Menzel S et al. 2014. HBS1L-MYB intergenic variants modulate fetal hemoglobin via long-range MYB enhancers. The Journal of clinical investigation 124: 1699-1710.

Steinberg MH, Chui DH, Dover GJ, Sebastiani P, Alsultan A. 2014. Fetal hemoglobin in sickle cell anemia: a glass half full? Blood 123: 481-485.

Teixeira SM, Cortellazzi LC, Grotto HZ. 2003. Effect of hydroxyurea on G gamma chain fetal hemoglobin synthesis by sickle-cell disease patients. Brazilian Journal of Medical and Biological Research 36: 1289-1292.

Terasawa T, Ogawa M, Porter P, Karam J. 1980. G gamma and A gamma globin-chain biosynthesis by adult and umbilical cord blood erythropoietic bursts and reticulocytes. Blood 56: 93-97.

Thein SL, Wainscoat JS, Sampietro M, Old JM, Cappellini D, Fiorelli G, Modell B, Weatherall DJ. 1987. Association of thalassaemia intermedia with a beta-globin gene haplotype. British Journal of Haematology 65: 367-373.

Uda M, Galanello R, Sanna S, Lettre G, Sankaran VG, Chen W, Usala G, Busonero F, Maschio A, Albai G et al. 2008. Genome-wide association study shows BCL11A associated with persistent fetal hemoglobin and amelioration of the phenotype of beta-thalassemia. Proceedings of the National Academy of Sciences of the United States of America 105: 1620-1625.

Vathipadiekal V, Alsultan A, Baltrusaitis K, Farrell JJ, Al Rubaish A, Al Muhanna F, Naserullah Z, Alsuliman A, Patra P, Milton JN et al. 2016. Homozygosity for a Haplotype in the HBG2-OR51B4 Region is Exclusive to Arab-Indian Haplotype Sickle Cell Anemia. American Journal of Hematology 91: E308-311.

Wood W. 2001. Hereditary Persistence of Fetal Hemoglobin and Thalassemia, in, Disorders of Hemoglobin: Genetics, Pathophysiology and Clinical Management, Chapter15, Steinberg MH, Forget BG, Higgs DR, Nagel RL eds. pp. 356-388. Cambridge University Press.

Xu J, Shao Z, Li D, Xie H, Kim W, Huang J, Taylor JE, Pinello L, Glass K, Jaffe JD et al. 2015. Developmental control of polycomb subunit composition by GATA factors mediates a switch to non-canonical functions. Molecular Cell 57: 304-316.

Zhou D, Liu K, Sun C-W, Pawlik KM, Townes TM. 2010. KLF1 regulates BCL11A expression and *γ*- to β-globin gene switching. Nature Genetics 42: 742.

